# *pnetr*: An R package for the PnET family of forest ecosystem models

**DOI:** 10.1101/2024.11.26.625449

**Authors:** Xiaojie Gao, Zaixing Zhou, Scott V. Ollinger, Jaclyn Hatala Matthes, Wenzhe Jiao, Jonathan R. Thompson

## Abstract

1. Ecosystem models offer a rigorous way to formalize scientific theories and are critical to evaluating complex interactions among ecological and biogeochemical processes. In addition to simulation and prediction, ecosystem models are a valuable tool for testing hypotheses about mechanisms and empirical findings because they reveal critical internal processes that are difficult to observe directly.
2. However, many ecosystem models are difficult to manage and apply by scientists who lack advanced computing skills due to complex model structures, lack of consistent documentation, and low-level programming implementation, which facilitates computing but reduces accessibility.
3. Here, we present the ‘*pnetr*’ R package, which is designed to provide an easy-to-manage ecosystem modeling framework and detailed documentation in both model structure and programming. The framework implements a family of widely used PnET (net photosynthesis, evapotranspiration) ecosystem models, which are relatively parsimonious but capture essential biogeochemical cycles of water, carbon, and nutrients. We chose the R programming language since it is familiar to many ecologists and has abundant statistical modeling resources. We showcase examples of model simulations and test the effects of phenology on carbon assimilation and wood production using data measured by the Environmental Measurement Station (EMS) eddy-covariance flux tower at Harvard Forest, MA.
4. We hope ‘pnetr’ can facilitate further development of ecological theory and increase the accessibility of ecosystem modeling and ecological forecasting.

## 1 Introduction

Ecosystem models integrate ecological processes to reflect our understanding of how ecosystems function in terms of the interactions between biology, chemistry, and physics (Bacmeister et al., 2020; Bonan et al., 2019; Geary et al., 2020; Middendorp et al., 2016; Scheller et al., 2010, 2012). Ecosystem models play a critical role in evaluating the interactions among multiple ecological processes (Liang et al., 2023; Thompson et al., 2011), investigating the causal effects of certain phenomena (Gustafson et al., 2017; Liang et al., 2018), and simulating potential future scenarios (De Bruijn et al., 2014; Prinn, 2013; Shifley et al., 2017). Complementary to bivariate statistical correlations, testing new hypotheses and scientific findings in ecosystem models allows scientists not only to directly compare multiple alternatives but also to investigate complex ecological feedbacks, interactions, and consequences. The development of ecological forecasting and model-data fusion techniques provides a powerful tool for hypothesis testing by predicting short- and long-term changes and aligning them with multi-source new observations (Dietze, 2017; Gettelman et al., 2022; Lewis et al., 2023). Because not all variables are easily observable, conducting ecological forecasting using ecosystem models provides an opportunity to link unobservable to observable processes and evaluate hypotheses in an integrated systematic view (Lewis et al., 2023; Liu et al., 2021).

Despite recent advances, testing new hypotheses in most ecosystem models is not a trivial task. Existing model frameworks have several limitations in the applications of hypothesis testing. First, for computational efficiency, existing ecosystem modeling frameworks are normally implemented in programming languages that are more widely used in software engineering such as C++, C#, and FORTRAN (Huang et al., 2020; Hurrell et al., 2013; Koven et al., 2020; Ma et al., 2022; Rastetter et al., 2022; Scheller et al., 2010). These implementations increase computing speed, but can also pose barriers for ecologists to modify model structures. Consequently, while many new scientific findings have led to suggestions for new or altered representations of certain ecological processes (Richardson et al., 2012; Stocker et al., 2019), implementing these suggestions is often far from straightforward. Collaborating with the corresponding software developers is one solution but it slows the iteration process due to communications and transferring materials. Second, many existing models lack clear and consistent documentation in both model structures and programming. Such models are most accessible to small user groups who already understand the implemented processes and do not require consistent documentation. Commonly, a process in the model will be updated but the documentation will not (Gustafson et al., 2023), which increases the difficulty for beginners and outsiders to catch up. Although the complexity of ecosystem models typically involves some preliminary knowledge, consistent documentation with version control can help simplify the difficulty and increase accessibility (Shifley et al., 2017). Third, due to their complex structures, many ecosystem modeling frameworks are difficult to manage by non-programmers. Ecosystem models are meant to be complex, however, aggregation and simplification are helpful in many scientific explorations. Parsimonious models can facilitate scientific hypothesis testing by simulating the essential ecological processes but removing the ones that are too complex and not directly related to the science questions. Ecologists can be more confident in modifying parsimonious models without worrying about making mistakes in other processes.

To facilitate hypothesis testing using ecosystem models, we developed the ‘*pnetr*’ ecosystem modeling framework using the statistical R language to implement the family of the PhotosyNthesis and EvapoTranspiration (PnET) ecosystem models. The PnET model family is a collection of parsimonious forest ecosystem models that have been actively developed since the 1990s (J. Aber et al., 1995; J. D. Aber & Federer, 1992; J. D. Aber et al., 1993, 1997). The original PnET model operates at a monthly time step, it was developed in Visual Basic and later updated to C++. Source code for some old versions or sub-models of the PnET family were also adapted to MATLAB and R, however, limited documentation is available. There is also a monthly time step sub-model of PnET being implemented as a succession of the LANDIS-II landscape simulation platform using C# (De Bruijn et al., 2014). The operation at the daily time step was recently added in the C++ version. Compared to previous implementations of PnET, here we provide a comprehensive but easy-to-manage modeling framework, as well as detailed algorithm documentation, for major sub-models in the PnET family. Our implementation supports both daily and monthly time steps in a consistent manner for all models. We chose the R statistical programming language to make the source code easy to modify and to better assist ecologists with the abundant statistical tools in the community for conducting scientific practices such as hypothesis testing, ecological forecasting, and model-data fusion.

## 2 PnET and ‘*pnetr*’

The PnET model family consists of a nested series of models simulating carbon, water, and nitrogen dynamics in forest ecosystems. These simple and lumped-parameter models are built on two principal relationships: 1) maximum photosynthetic rate is a function of foliar nitrogen concentration, and 2) transpiration is a function of realized photosynthetic rate (J. Aber et al., 1995; J. D. Aber & Federer, 1992; J. D. Aber et al., 1993, 1997). The PnET models have been successfully used to predict gross and net primary productivity (GPP and NPP), carbon and water balances, and nitrogen dynamics in forest ecosystems responding to changes in climate, nitrogen deposition, land use, and species composition at both site and grid levels (De Bruijn et al., 2014; Liang et al., 2023; Ollinger et al., 1998; Zhou et al., 2018).

Our ‘*pnetr*’ R package provides three major PnET sub-models including PnET-Day (J. D. Aber et al., 1996), PnET-II (J. Aber et al., 1995; J. D. Aber & Federer, 1992), and PnET-CN (J. D. Aber et al., 1997). The sub-models differ in structures and perspectives but share common routines (Fig. 1). PnET-Day is the simplest model in the family, it only simulates the photosynthetic processes at leaf and canopy scales. PnET-II includes routines concerning water and carbon cycles, and PnET-CN further includes nitrogen cycles with complete feedbacks between carbon, water and nitrogen. We also provide tools for sensitivity analysis so that users can investigate the relationships between multiple variables when they modify the model structures.

**Figure 1.**
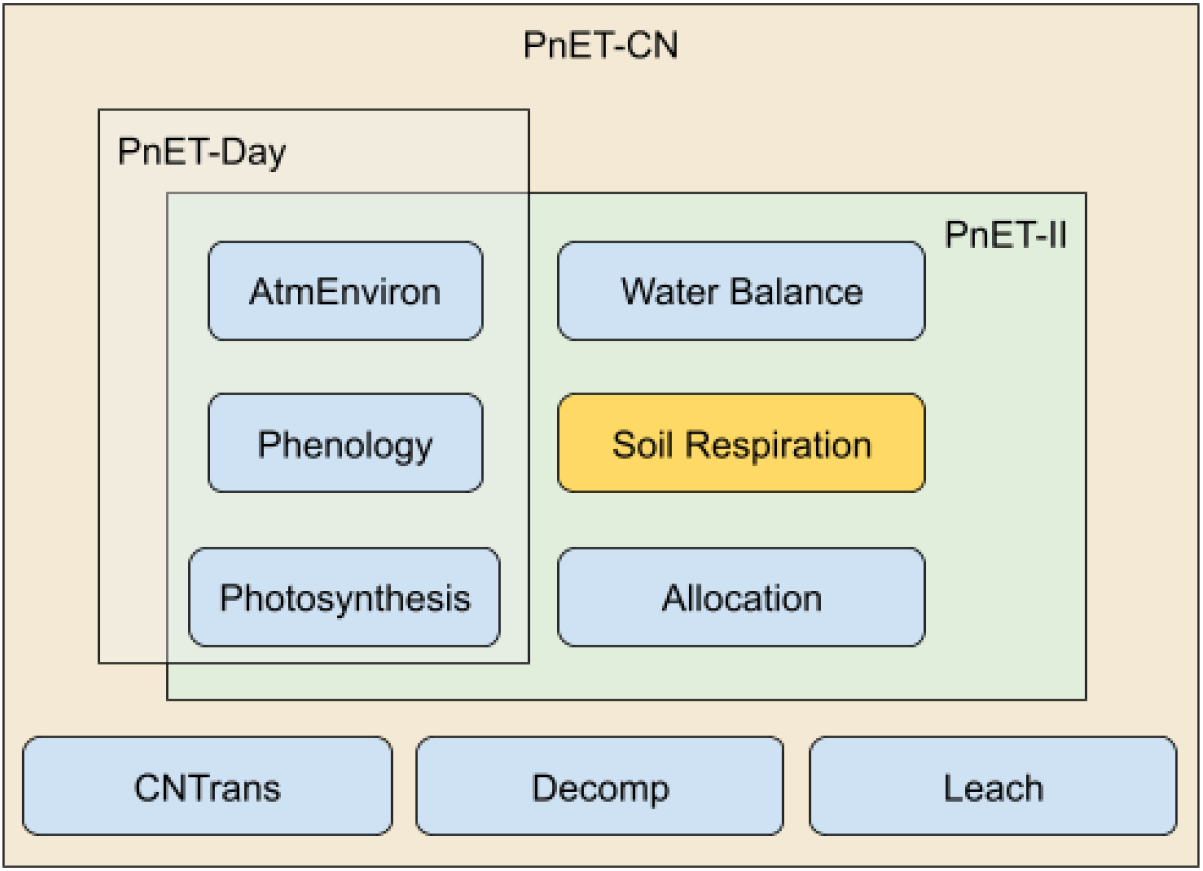
Major PnET sub-models and their components. Note that ‘Soil Respiration’ is included in the ‘Decomp’ routine when using PnET-CN.

There are nine major routines in the PnET models (Fig. 1). **’AtmEnviron’** deals with the atmospheric environment and calculates meteorological variables such as average temperatures and vapor pressure deficit needed to run the simulations. **’Phenology’** controls the timing of plants’ critical life-cycle events such as leaf development, woody growth, leaf senescence, and dormancy. **’Photosynthesis’** concerns the processes in carbon assimilation through photosynthesis, which are driven by foliar nitrogen concentration and affected by phenology and environmental factors such as light radiation, temperature, and water stress. **’Water Balance’** simulates the water cycle in the field including precipitation, snow, and evapotranspiration. **’Soil Respiration’** quantifies the amount of carbon emitted from respiration in soil. **’Allocation’** is the process that allocates the assimilated carbon and/or nitrogen into different pools in the plant including the whole plant pool, the bud pool, the wood pool, and the root pool. Allocation only occurs monthly and annually. **’CNTrans’, ‘Decomp’**, and **’Leach’** determine the nitrogen cycle, which dynamically computes foliar nitrogen concentration over the years. The nitrogen cycle includes nitrogen deposition and litterfall to soil organic matter transfer, nitrogen mineralization and nitrification, and leaching losses of nitrate. The effects of CO_2_ fertilization (Ollinger et al., 2002) and Ozone concentration (Ollinger et al., 1997) on photosynthesis are also included in the PnET-CN model.

A simple schematic diagram of the PnET-CN model illustrates the major processes included in the PnET models (Fig. 2). During the growing season, carbon is fixed through leaf photosynthesis, quantified by gross primary productivity (GPP), and released by foliar respiration at the same time. The assimilated carbon is then transported to the plant carbon mobile pool and allocated to grow leaves, wood, and roots after deduction of autotrophic respiration. This net carbon storage is represented by net primary productivity (NPP). Carbon emissions from the decay of plant detritus and decomposition of soil organic matter are also quantified. The rate of photosynthesis depends on the nitrogen concentration in the leaves and is affected by environmental factors including light radiation, temperature, vapor pressure deficit, and water stress. The only source of water is precipitation. Depending on the temperature and site conditions, water entering the system can be in the form of rain and/or snow. The amount of water available for plant uptake is affected by processes such as fast flow and drainage. Plants lose water through evapotranspiration. The nitrogen cycle is coupled with the carbon cycle. The foliar nitrogen concentration determines the maximum net photosynthesis rate without water stress, which increases the internal non-structure plant carbon pool and thus NPP. Conversely, the increased plant NPP will increase the nitrogen demand, which reduces the non-structural plant nitrogen pool. The degree to which nitrification occurs is negatively correlated with the strength of plant demand for nitrogen in competition with nitrifiers. The amount of NO3^-^ and NH4^+^ in the soil solution affects plant nitrogen uptake and the nitrate leaching rate.

**Figure 2.**
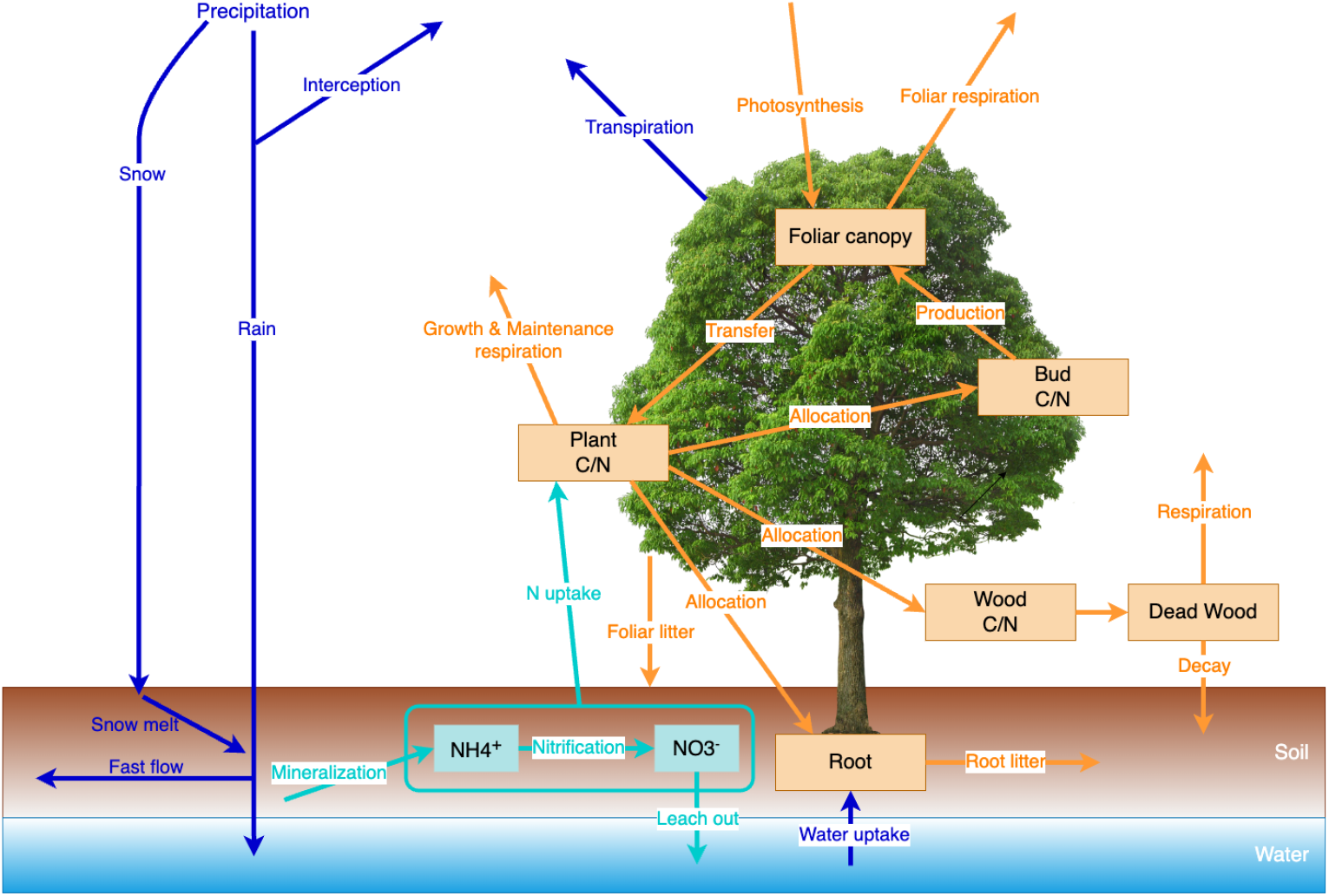
PnET-CN schematic diagram. Arrows in color represent the major processes in the biogeochemical cycles including carbon (yellow), water (blue), and nitrogen (cyan).

## 3 Working Examples

Here we use three working examples including carbon dynamics simulation, sensitivity analysis, and hypothesis testing to demonstrate the general usage of the ‘pnetr’ R package and how it can facilitate scientific exploration.

### 3.1 Simulating Ecosystem Photosynthesis

To run the model using the ‘pnetr’ package, users need input data describing climate conditions, and site and vegetation characteristics (Table 1). The required climate data vary by sub-models, but they generally include maximum (Tmax) and minimum temperature (Tmin), photosynthetic active radiation (PAR), precipitation, CO_2_ and O_3_ concentration in the air, and NO_3_ and NH_4_ deposition. The climate data can be daily, weekly, or monthly, and its resolution determines the time-step of the model simulation. The site information includes the latitudinal location, soil water holding capacity, and initial snowpack. The vegetation characteristics include a series of empirical values indicating information such as foliar nitrogen concentration, maximum foliar mass, phenological requirements, respiration rate, etc. Default values for most vegetation characteristics are provided. The full list of the input data can be found in the package documentation.

**Table 1.**
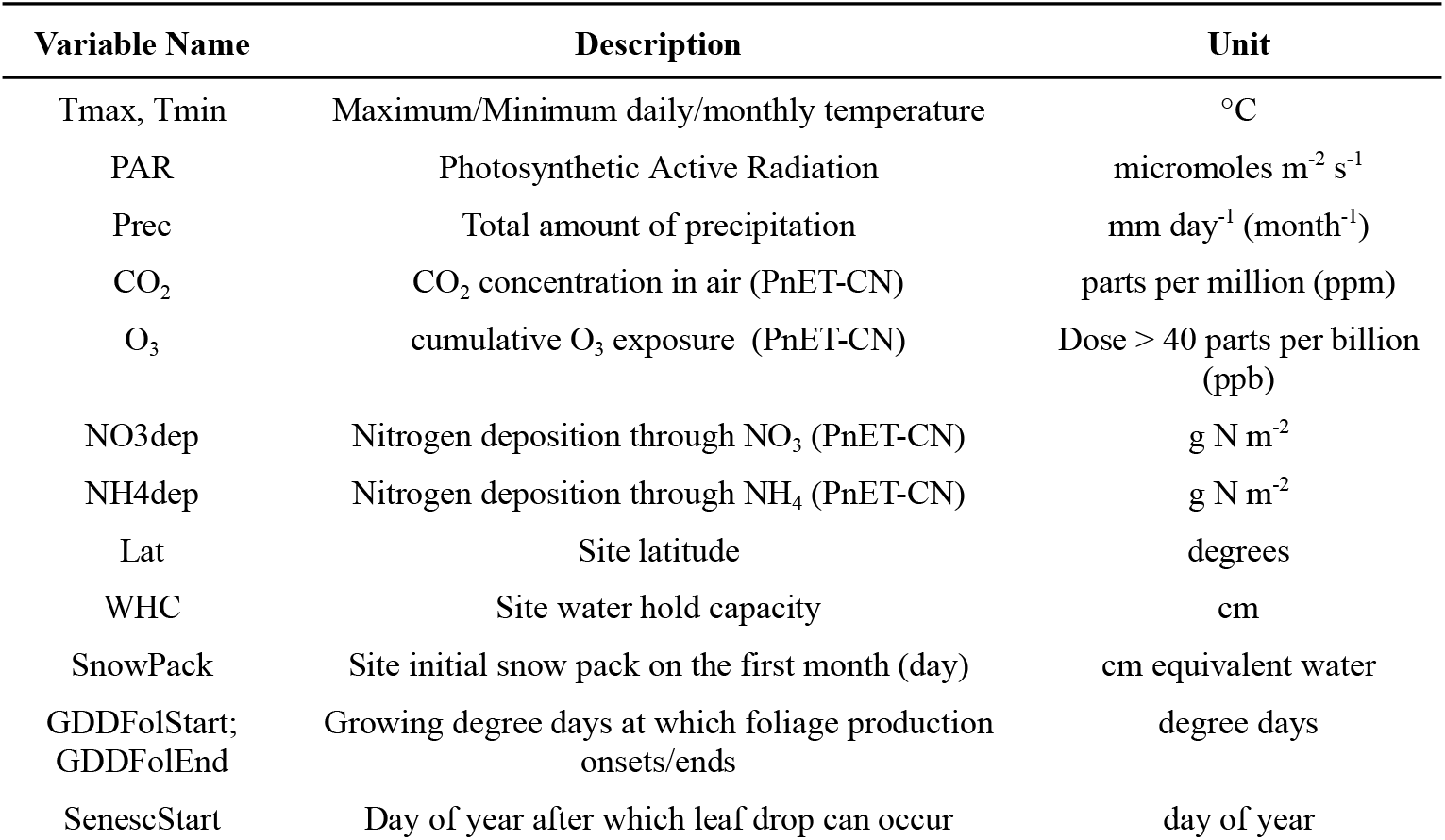
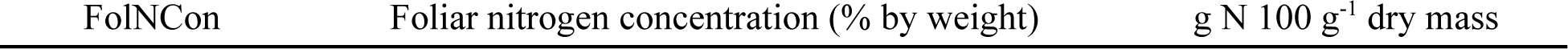
Some important input parameters for PnET model simulations (The full list and the corresponding default values can be found in the ‘pnetr’ package documentation).

As an example, we simulated gross primary productivity (GPP), net ecosystem CO_2_ exchange (NEE), and plant internal carbon pools and evaluated model performance using data collected by the Environmental Measurement Station (EMS) eddy-covariance flux tower at the Harvard Forest, MA (Munger & Wofsy, 2024). Air temperature input variables (Tmax, Tmin) and incoming photosynthetic radiation flux (PAR) were aggregated to daily values from hourly measurements at the Harvard Forest long-term meteorological station (Boose et al., 2024). The EMS tower measured hourly net ecosystem exchange (NEE; *i*.*e*., the net ecosystem-atmosphere carbon flux) over the 1992-2023 time period. The EMS NEE data were filtered to remove periods of low turbulence using a seasonally variable friction velocity threshold (Pastorello et al., 2020) where the growing and non-growing seasons were defined by changes in Landsat-derived leaf greenness seasonality within the tower footprint (Gao et al 2021). EMS tower GPP was partitioned from NEE by fitting a temperature-light response function to NEE data (Munger & Wofsy, 2024). Tower GPP and NEE were aggregated to daily and monthly sums for comparison to the PnET model output. Tower NEE was used to compare with PnET-modeled NEE.

We ran all PnET sub-models at both monthly (Fig. 3, Fig. 4) and daily steps (Fig. 5). In general, model simulations are reasonably consistent with eddy-covariance measurements. The simulated GPP values in PnET-Day and PnET-II are similar due to limited water stress at this site. PnET-CN simulated GPP values align better with the EC measurements, suggesting the importance of accounting for the nitrogen cycle effect on photosynthesis. Although all models underestimate the higher GPP and NPP values during 2002-2012, their general variability is fitted reasonably well, especially for GPP. The dynamics of carbon in the plant components show the carbon storage and allocation strategies for above and below-ground growth (Fig. 4).

**Figure 3.**
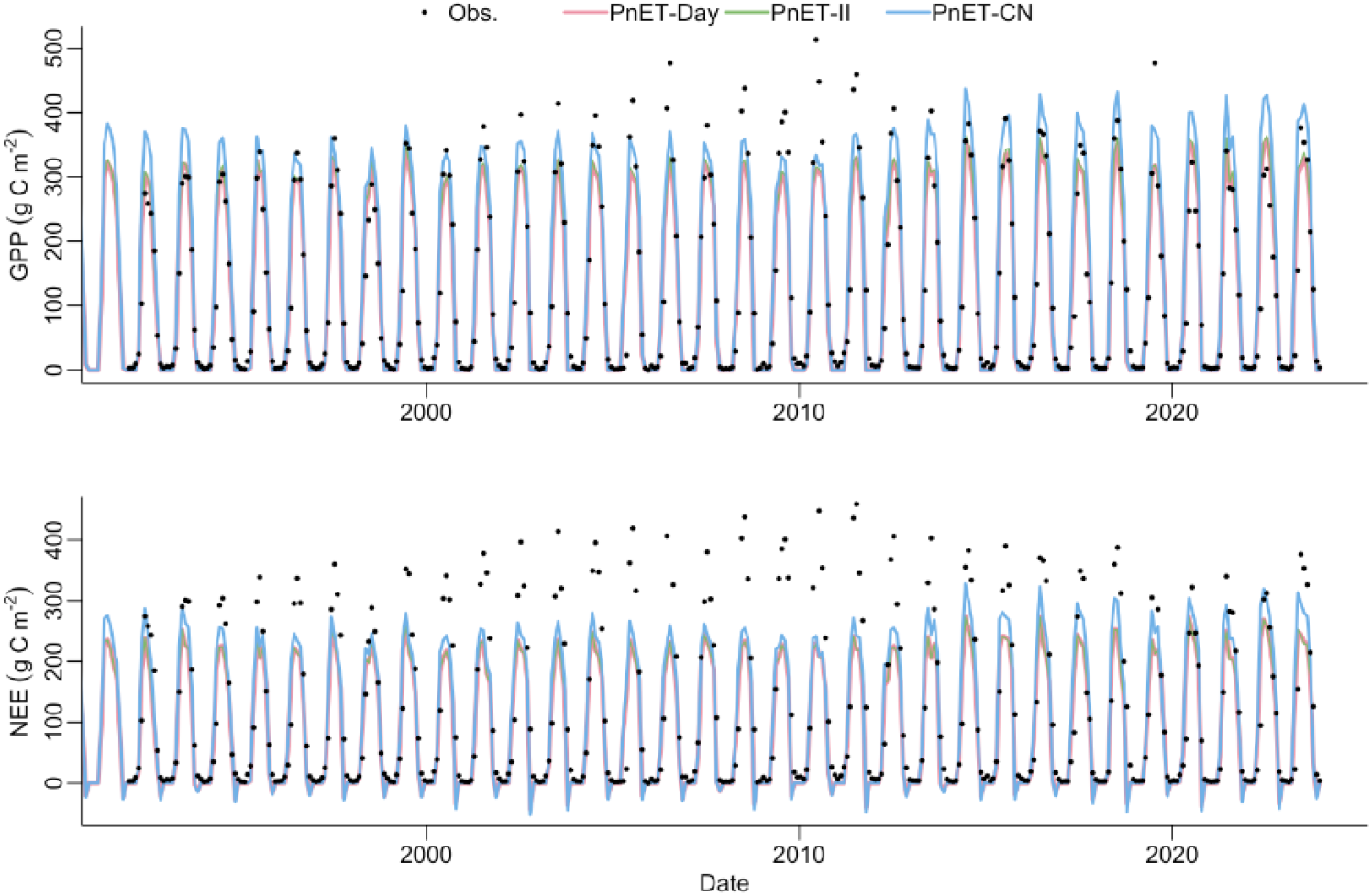
PnET models simulated GPP and NEE compared with the measured data collected by the Environmental Measurement Station (EMS) eddy-covariance flux tower installed at Harvard Forest, MA. Models were simulated in monthly steps.

**Figure 4.**
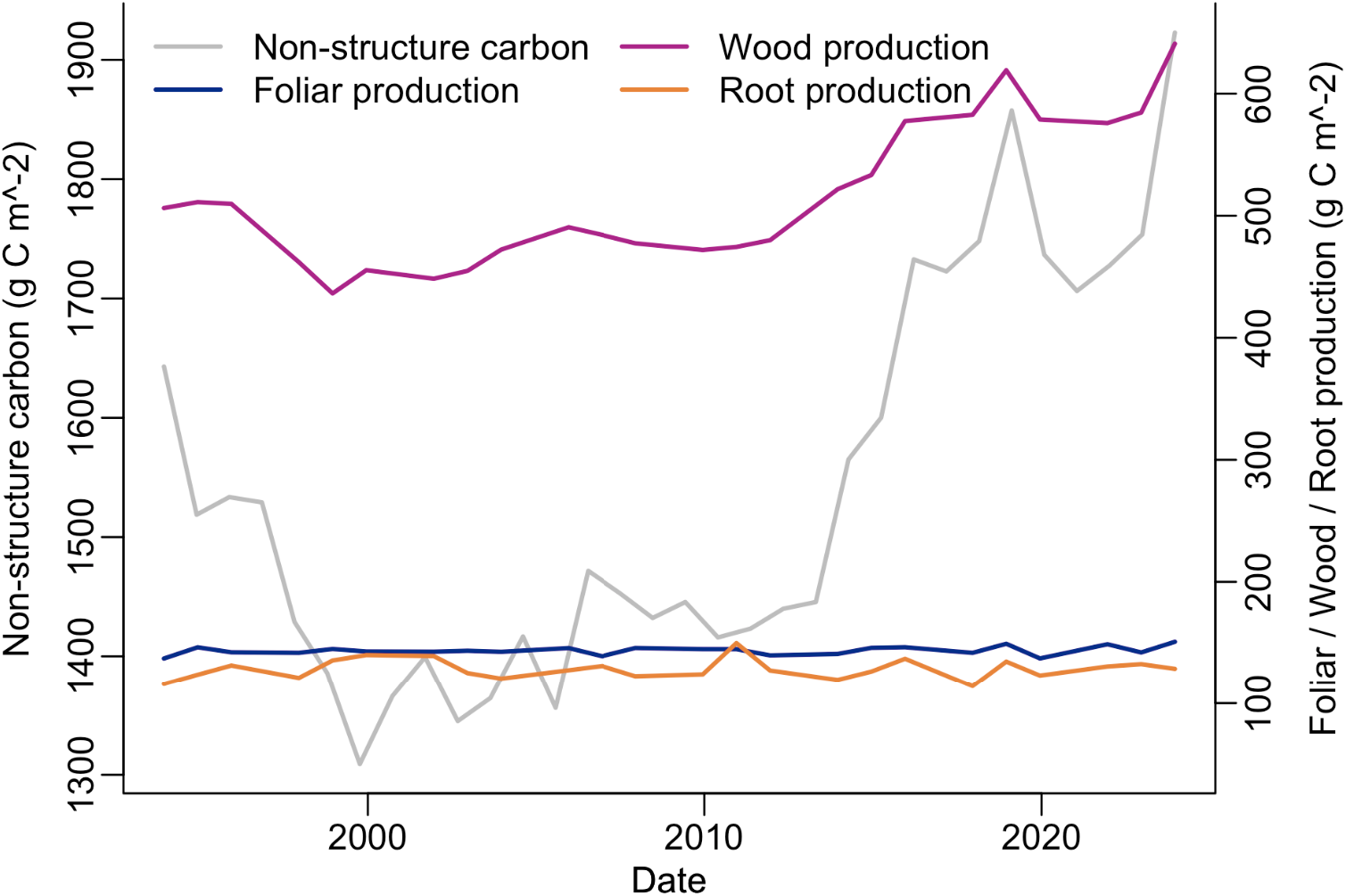
Annual non-structural carbon dynamics and net primary production simulated by the PnET-CN model. Annual net primary production is allocated to store carbon in foliar, wood, and root production.

**Figure 5.**
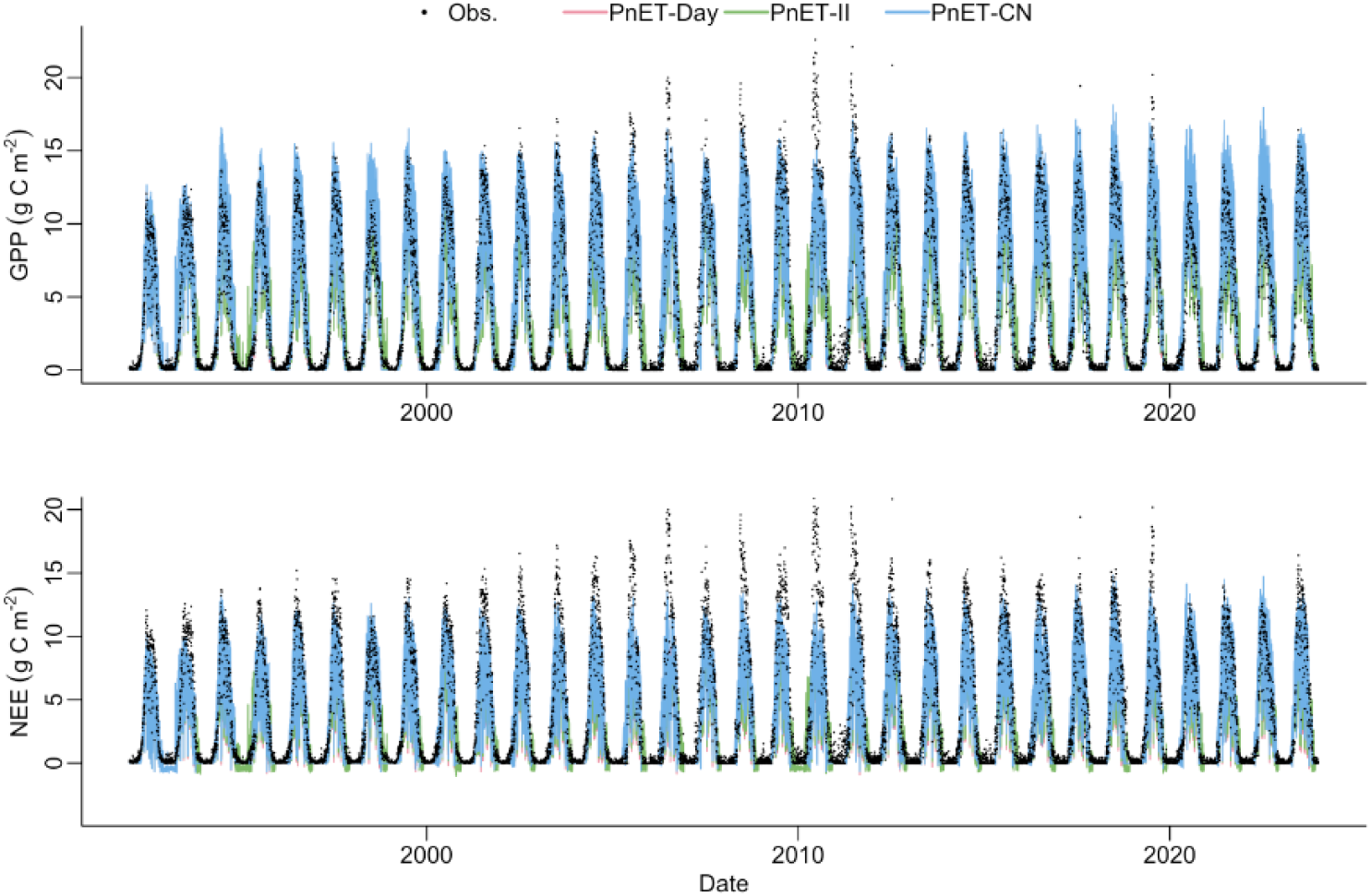
PnET models simulated GPP and NEP compared with the measured data collected by the Environmental Measurement Station (EMS) eddy-covariance flux tower installed at Harvard Forest, MA. Models were simulated in daily steps.

The daily simulations reveal similar patterns to the monthly simulations but with more pronounced temporal variations (Fig. 5). The daily versions of models enable investigating processes operating on shorter temporal scales (e.g., the daily variation of photosynthesis and water balances). Similar to the monthly results, although PnET models reasonably simulated the variations of carbon exchanges at this site, the consistent underestimation in the summer suggests that there is potential for improvements and calibration in model structures and parameter values. Indeed, this underestimation of the summer GPP problem at some sites has been noticed (Zhou et al., 2018), although remains challenging. The ‘pnetr’ R package makes the modeling framework transparent and increases accessibility, which will better assist ecologists in testing new theories and improving the simulations. We revisit this issue in Section 3.3 on hypothesis testing.

### 3.2 Sensitivity Analysis

Sensitivity analysis is often conducted to investigate the sensitivity of model output variables to input and/or internal parameter variations (McKenzie et al., 2019). We encourage users to modify the processes implemented in the ‘*pnetr*’ package for their purposes, so ‘pnetr’ includes a routine for users to easily conduct sensitivity analysis. First, a simple automatic sensitivity analysis can be performed by calling an integrated function. In this way, parameter variations are prescribed by varying their default values by 10% using a Monte Carlo Markov Chain (MCMC) sampling procedure. Note that caution needs to be taken using this method with user-defined parameter values, especially for monthly simulations, as slight changes in some parameters may be exaggerated in monthly scales. For instance, when running the models at a monthly step, the unit of variables controlling phenology (e.g., start-of-season, start-of-senescence) is a month below even though phenological changes may be in several days, it could result in a different month, leading to a large effect on the model simulation. The automatic sample generation can be used as a template for users to modify as needed. Alternatively, a manual parameter sampling procedure in which the distributions of parameters can be defined by the users. This method is a bit tedious but gives full control to the users. Second, after obtaining the parameter samples, we run the model with each parameter set and collect the user-specified target output variables (e.g., GPP). Parallel processing can be utilized in this step to reduce computing time. Finally, we perform a random forest regression to evaluate and visualize the importance of each parameter in explaining the target output variables.

As an example, here we show annual GPP and NPP sensitivity to 10% parameter variations in the PnET-CN model running at a monthly step (Fig. 5). The meanings of the parameters can be found in the package documentation. More importantly, note that this is just a toy example and may provide limited information about the model processes. Some parameters showing large sensitivity, such as the constant used to calculate water-use efficiency (WUEconst) from vapor pressure deficit (VPD), the slope of the relationship between foliar N and max photosynthetic rate (AmaxB), and the fraction of carbon in foliage mass (CFracBiomass) showed large control to GPP and/or NEP in the example. However, these parameters are usually prescribed using empirical values from observational analysis, so they typically produce less variability in model predictions than in an unconstrained sensitivity analysis. Therefore, this example is to show the sensitivity analysis feature of the package, to acquire useful insights about model processes, users should carefully determine which parameters are of interest and which parameters are viewed as constants.

### 3.3 Hypothesis Testing with ‘pnetr’

Plant phenology is the first-order control of carbon exchange in vegetated ecosystems. Previous studies show that earlier plant spring leaf-out can significantly increase annual GPP, which measures the total amount of carbon absorbed (Gao et al., 2023; Keenan et al., 2014). However, earlier springs may not always increase wood production depending on many complex processes in carbon sequestration and allocation (Dow et al., 2022). Here, as a simple example, we use the daily PnET-CN model to test the hypothesis that spring phenology is not significantly correlated with wood production.

Specifically, we modified the phenology routine of the PnET-CN model to include historical spring and autumn phenology observations and quantified the relationship between the model simulated annual GPP and historical spring phenology. The historical phenology information in 1984-2023 was obtained from Landsat observations using a Bayesian land surface phenology model (BLSP; Gao et al., 2021). To align with the PnET-CN model’s phenology routine, we used a threshold-based method to transform BLSP phenometrics as growing-degree-day-controlled dates of start-of-foliar-development (GDDFolStart; Table 1) and start-of-senescence (SenescStart; Table 1) (Fig. 7a). We then substituted both spring and autumn phenology in the model with BLSP observations and investigated the impacts of historical changes in spring phenology on annual GPP, plant carbon pool (PlantC), and wood production (WoodProdCYr), respectively. The R implementation of the ‘pnetr’ package makes model modification and visualization simple.

**Figure 6.**
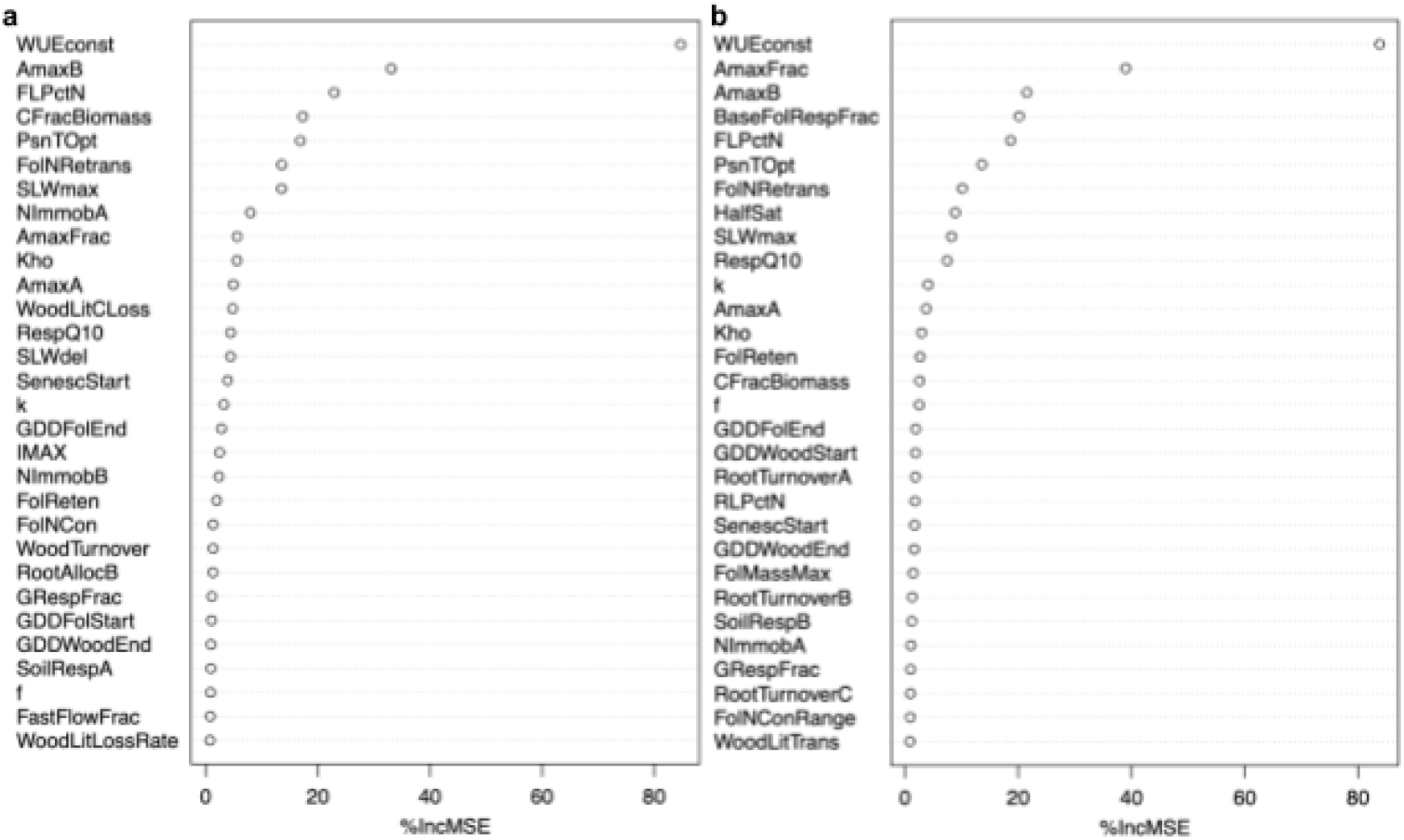
Variable importance of the monthly PnET-CN model in simulating annual GPP (a) and NEP values (b). The sensitivity analysis was iterated 1000 times with each time randomly changing the default values of the variables by 10%. Variable names and descriptions can be found in the ‘pnetr’ package documentation.

**Figure 7.**
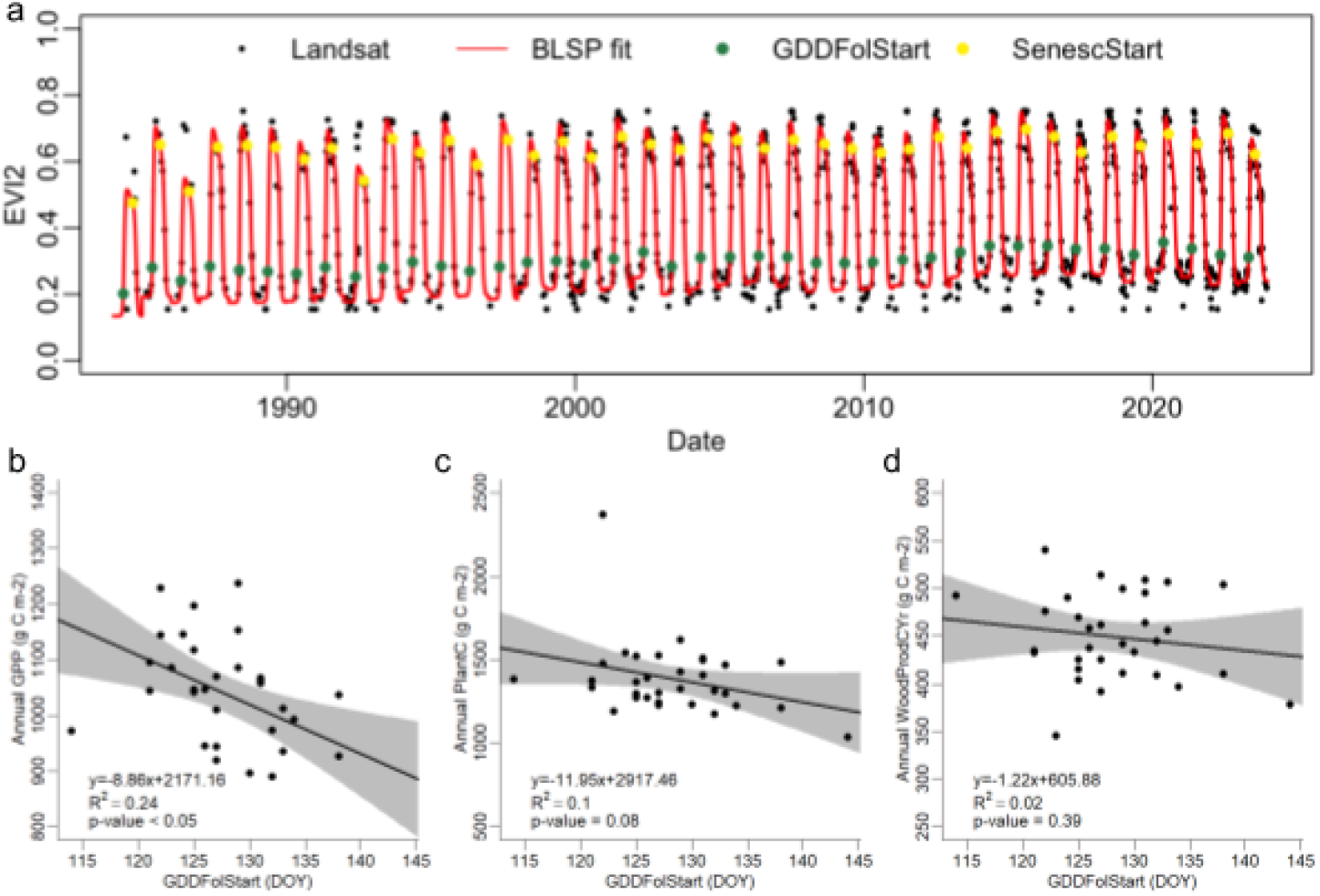
Testing the effects of spring and autumn phenology on carbon exchange. Note that ‘GDDFolStart’ and ‘SenescStart’ are PnET model parameters controlling for the dates of the start of foliar development and the start of senescence, here we substitute these dates with satellite observed phenological dates, thus their units are day-of-year (DOY) in the figure. Phenology information was retrieved from the fitted time series of Landsat observations at Harvard Forest Environmental Measurement Station (EMS) eddy-covariance flux tower site using a Bayesian land surface phenology (BLSP; Gao et al 2021) model (a). To match with the PnET-CN model parameters, we use the time when the EVI2 trajectory reaches 15% of its amplitude as the threshold to determine ‘GDDFolStart’ DOY and 95% of its amplitude to determine ‘SenescStart’ DOY. The relationships between spring phenology and annual gross primary productivity (GPP; b), plant carbon pool (PlantC; b), and wood production (WoodProdCYr; d) are shown in scatter plots, in which the linear regression lines, confidence intervals (grey polygons), and statistical summary of the model fits are also shown.

The simulation result shows that earlier spring phenology is significantly correlated with higher annual GPP (Fig. 7a; R^2^ = 0.24; p-value < 0.05) and weakly correlated with higher annual PlantC (Fig. 7b; R^2^ = 0.10; p-value = 0.08) but not with WoodProdCYr (Fig. 7c; R^2^ = 0.02; p-value = 0.39), which suggests that spring phenology significantly affects carbon sequestration but may not directly affect wood production due to complex internal carbon processes. This result is consistent with previous studies (Dow et al., 2022; Keenan et al., 2014), although the simulation is a relatively simple example and the extension of the conclusion needs more investigation. More importantly, however, this simulation example shows that testing hypotheses in ecosystem models can reveal complex interactions between multiple variables and suggest how the apparent relationships propagate internally. Thus, the result of the model simulation can be used as either proof of observational phenomena or as clues for further investigations.

## 4 Applications

We envision four major applications of the ‘pnetr’ R package. First, model simulations. This is the most direct application. Once the parameters are carefully prescribed at a site, the models can simulate carbon, water, and nitrogen cycles to better understand the biogeochemical processes. Using projected future environmental variables, model simulation provides plausible future scenarios, which can be a useful tool to understand future climate change impacts. Second, hypothesis testing. As we have shown, hypothesis testing is straightforward and it allows investigations of interactions of multiple ecological processes. In addition to our example, the daily scale PnET-CN model has been used in evaluating how different process-based spring phenology models representing different hypotheses affect the simulations of photosynthetic productivity (Teets et al., 2023). The model’s embedded processes represent our current ecological theory, and if a new scientific finding can consistently increase the accuracy of model simulations, it is a direct sign of theory advancement. Third, model-data fusion. Since neither the model nor the data are perfect, model-data fusion using Bayesian methods is a useful technique to integrate various big data into model simulations and quantify uncertainty. Model-data fusion can also be used to retrieve internal variables that are difficult to measure (Liu et al., 2021), or to develop near-real-time ecological monitoring systems (Dietze, 2017; Fer et al., 2018). Our R implementation benefits ecologists with abundant statistical resources, especially Bayesian methods, which are critical to model-data fusion. Fourth, hands-on education. Although the package was developed mainly for scientific exploration, it can also be used as a tool for students to learn biogeochemical processes and get hands-on experience in ecological modeling. The elaborate documentation we provide can help students understand the variables and algorithms in ecology.

## 5 Conclusion

With the aim of providing an easy-to-manage ecosystem modeling framework that captures essential carbon, water, and nutrient processes for ecologists, we developed the ‘pnetr’ package to implement a family of PnET ecosystem models using the R programming language. We provide detailed documentation about the implemented algorithms to help users better understand the models and be confident in modifying the processes to suit their needs. Compared to more complex ecosystem models, the ‘pnetr’ package helps scientists focus more on ecology without the burden of acquiring savvy skills in computer science. Additionally, the R implementation gives users easy access to abundant statistical resources for modeling and visualization. Here, we present the package, provide working examples, and propose applications. We hope the ‘pnetr’ R package can facilitate ecological theory development and scientific hypothesis testing, and increase the accessibility of ecosystem modeling.

## Acknowledgements

X.G. and J.R.T. are funded by NSF-DEB-LTER Grant 18-32210 and NSF-DISES Grant 22-05705. J.H.M. was funded by NSF-DEB Grant 22-31681 and NSF-EF Grant 19-26454. Z.Z. and S.V.O. were funded by NSF-EPSCoR Grant 1920908, NSF-DEB-LTER Grants 1832210, and the New Hampshire Agricultural Experiment Station, Hatch project NH00711.

## Author contributions

X.G. and J.R.T conceptualized the idea. X.G. developed the package, documented the algorithms, and conducted the analyses with important feedback from J.R.T., Z.Z., S.V.O., J.H.M, and W.J.; J.H.M. formatted the long-term eddy-covariance measurements for the model tests. X.G. drafted the manuscript and all authors contributed to editing and revising. All authors approved the final manuscript.

## Code and data availability

The R package is available on GitHub (https://github.com/hf-thompson-lab/pnetr). An example dataset to run the model is included in the package repository. The dataset was derived from the data measured by the Environmental Measurement Station (EMS) eddy-covariance flux tower at Harvard Forest, MA (https://harvardforest.fas.harvard.edu/research/towers) and the FLUXNET2015 dataset (https://fluxnet.org/data/fluxnet2015-dataset, site ID: US-Ha1).

